# Sequence-specific model for predicting peptide collision cross-section values in proteomic ion mobility spectrometry

**DOI:** 10.1101/2020.09.14.296590

**Authors:** Chih-Hsiang Chang, Darien Yeung, Victor Spicer, Oleg Krokhin, Yasushi Ishihama

## Abstract

The contribution of peptide amino-acid sequence to collision cross-section values (CCS) has been investigated using a dataset of ∼134,000 peptides of four different charge states (1+ to 4+). The migration data was acquired using a two-dimensional LC/trapped ion mobility spectrometry/quadrupole/time-of-flight MS analysis of HeLa cell digests created using 7 different proteases and was converted to CCS values. Following the previously reported modeling approaches using intrinsic size parameters (ISP), we extended this methodology to encode the position of individual residues within a peptide sequence. A generalized prediction model was built by dividing the dataset into 8 groups (four charges for both tryptic/non-tryptic peptides). Position dependent ISPs were independently optimized for the eight subsets of peptides, resulting in prediction accuracy of ∼0.981 for the entire population of peptides. We find that ion mobility is strongly affected by the peptide’s ability to solvate the positively charged sites. Internal positioning of polar residues and proline leads to decreased CCS values as they improve charge solvation; conversely, this ability decreases with increasing peptide charge due to electrostatic repulsion. Furthermore, higher helical propensity and peptide hydrophobicity result in preferential formation of extended structures with higher than predicted CCS values. Finally, acidic/basic residues exhibit position dependent ISP behaviour consistent with electrostatic interaction with the peptide macro-dipole, which affects the peptide helicity.

## INTRODUCTION

Ion mobility spectrometry (IMS) has been long considered as a promising tool for many applications in structural biology,^1^ proteomics^2^ and many other analytical applications.^3,4^ Separation of isobaric peptides,^5^ improving signal-to-noise ratio in bottom-up approaches,^6^ studying protein conformation and protein assemblies^7^ represents an incomplete list of its proteomic applications. One of the attractive options of IMS is the possibility to model gas-phase peptide behaviour in ion-mobility based separation. Building a comprehensive collisional cross section (CCS) prediction model for peptides will allow not only the direct application to improve confidence of MS/MS-based identification^8^ but will help better the current understanding of underlying mechanisms for ion mobility-based separations, resulting in improving MS/MS-based quantitation by reducing the complexity of peptide ions prior to tandem mass spectrometry.^9,10^

Considering the history in the progress of this field, it is easy to notice striking similarity between IMS and reversed-phase high performance liquid chromatography (RP-HPLC). Both were conceived long before arrival of MS based proteomics.^11-13^ Both techniques are used as front-end devices to improve delivery of separated compounds into the mass spectrometer and are amenable to modeling of the separation processes – driven by peptides’ size/shape in the gas phase and hydrophobicity, respectively.^8,14^

Due to its preparative capability, RP-HPLC became one of the most important techniques in protein/peptide analytical chemistry long before the proteomic era. Initial peptide retention time prediction models aimed at improving separation method development during early 1980s.^14,15^ These early models were based on an additive principle, which considers that the hydrophobicity for a peptide is equal the sum of its constituent residues’ hydrophobicities. The effect of peptide sequence in addition to its composition in RP HPLC has been reported in 1987^16^ and first sequence specific model has been developed in 2004 based on the collection of just 346 tryptic peptides.^17^ The authors suggested using position-dependent hydrophobicity coefficients for individual amino acids to compensate for unique features of peptide N-termini observed due to ion-pairing interactions; however, sufficient data density is required to model this concept. Development of mass spectrometry and proteomics brought increased throughput and confidence in peptide identifications, thus increasing the size of high-quality datasets available for prediction modeling.^18^ Proteomic peptide datasets have allowed the implementation of more complex prediction algorithms and opportunities to study a variety of structural features in peptides. In mid 2000s, Petritis et al. described retention modeling via an artificial neural network (ANN) approach using datasets of ∼6,000^19^ and later ∼300,000 peptides.^20^ The increasing size of proteomic datasets near the 2010s opened the opportunity to study the effects of structural motifs such as N-cap helical stabilization on peptide retention in RP HPLC. Given only a small portion of peptides exhibit amphipathic helicity, such study required a collection of ∼280,000 peptides.^21^ Continuous efforts in standardization of RP HPLC separation in proteomics^22,23^ and progressive growth in MS productivity in the past decade has allowed for the collection of high quality RP HPLC data in the size of hundreds of thousand to over a million peptides.^24-27^ This paved the possibility for wider application of high data density machine learning techniques to address peptide retention time prediction problems.^27-29^

Clemmer and co-workers led the way in the development of IMS technology for proteomic applications,^30^ peptide IMS data collection, and modeling peptide ion mobility.^8,31^ Valentine et al. used 660 peptides to derive the intrinsic size parameter (ISP) coefficients, which multiplicatively scales the mass of individual residues used in CCS additive prediction models.^31^ The same group of authors expanded this algorithm to the collection of 2,094 tryptic peptides 5-15 amino acids long.^8^ In a different approach, Shah et al. built a machine-learning based model attempting to introduce additional features including but not limited to: normalized retention time in RP HPLC, peptide length, gas phase basicity, and number of negatively/positively charged groups.^32^ However, the size of this dataset, which contained 3933 (2+), 3916 (3+), 717 (4+ peptides), was not sufficient to define sequence-specific features. Peptide structural properties are of ultimate importance for IMS. The same peptide species can assume different conformations with drastically different CCS values.^33^ This feature is the most obvious for 3+ peptide populations, which exhibit significant split in CCS versus molecular weight plots – designated as compact and extended structure populations.^32,34,35^ Another argument confirming sequence dependent character of peptide IMS was provided by Lietz et al.^34^ The authors used Lys-C and Lys-N digests of K562 to show that N-terminal location of Lys results in lower CCS values compared to the same sequences with C-terminal Lys for 14 peptide pairs.

Overall, there is ample evidence of sequence-dependent character of IMS separation. Yet, compared to RP HPLC, there are no CCS prediction models incorporating these features. One of the problems is a significant advantage of chromatographic separations in terms of data availability: hundreds of thousand data points^23-25,27^ vs. thousands.^8,32^ The timsTOF Pro, a quadrupole/time-of-flight (Q/TOF) mass spectrometer coupled with trapped ion mobility spectrometry (TIMS) cells, achieves a resolving power of over 220K and the scan speed (100 ms per scan) between LC and Q/TOF mass analyzer, showed a lot of promise in this regard.^35-37^ Similar to chromatographic applications, measuring CCS values for few hundred thousand peptides may provide sufficient data for application of machine learning approaches. At the time of our work, Meier et al. concurrently developed a deep learning CCS prediction model using 570,000 unique combinations of sequence, charge state (2+, 3+ and 4+), including peptides with oxidized methionine.^38^ However, many machine learning approaches often operate as “black boxes”, providing limited information on the underlying separation mechanisms. Meier et al. have demonstrated the contributions of the 20 amino acid residues and the qualitative trends for hydrophobicity, Pro content, and position of His.^38^ They highlighted the difficulty to model the observed physicochemical properties along with sequence dependent features directly with the linear sequences and our work here is able to address such difficulties as well as investigate finer composition and position dependent features that are novel to our approach. Therefore, a semi-empirical Sequence-Specific Retention Calculator (SSRCalc) approach based on position-dependent correction coefficients was applied in this work to establish a Sequence-Specific Ion mobility Calculator (SSICalc).^39^ The SSRCalc has been used successfully in the past for modeling various modes of peptide HPLC,^39-41^ and capillary zone electrophoresis.^42^ The dataset for CCS modeling was obtained by the 2D LC (SCX/RP)/ESI/TIMS/Q/TOF analysis of multiple alternative proteases digests using timsTOF Pro.

## EXPERIMENTAL SECTION

### Materials

Ammonium bicarbonate (ABC), 2-amino-2-(hydroxymethyl)-1,3-propanediol hydrochloride (Tris-HCl), sodium deoxycholate (SDC), ammonium acetate (AA), sodium N-lauroylsarcosinate (SLS), tris(2-carboxyethyl)phosphine (TCEP), 2-chloroacetamide (CAA), calcium chloride, ethyl acetate, acetonitrile, acetic acid, trifluoroacetic acid, V8 protease (Glu-C), lysyl endopeptidase (Lys-C) and other chemicals were purchased from Fujifilm Wako (Osaka, Japan). Modified trypsin, chymotrypsin and Asp-N / Lys-N / LysargiNase were procured from Promega (Madison, WI) / Thermo Fisher Scientific (Waltham, MA) / Merck (Darmstadt, Germany), respectively. Polystyrene-divinylbenzene (SDB) and cation exchange-SR (SCX) Empore™ disks were purchased from GL Sciences (Tokyo, Japan). Water was purified by a Millipore Milli-Q system (Bedford, MA).

### HeLa cell culture and protein extraction

HeLa S3 (human cervical adenocarcinoma) cells were cultured to 80% confluence in 10-cm diameter dishes then harvested in lysis buffer containing protease inhibitors (Sigma-Aldrich, St. Louis, MO), 12 mM SDC, 12 mM SLS, 10 mM TCEP, 40 mM CAA in 100 mM Tris buffer (pH 8.5). The lysate was vortexed and sonicated on ice for 20 min. The final protein concentration of the sample was determined using the bicinchoninic acid (BCA) protein assay (Thermo Fisher Scientific).

### Protein Digestion

The proteins were digested using previously described phase-transfer surfactants (PTS) method.^43^ For LysargiNase digestion, protein extract was diluted 10-fold by using 10 mM CaCl_2_ and digested with LysargiNase (1: 40 w/w) overnight at 37 °C. For other proteases, extracts were diluted 5-fold by with 50 mM ABC and digested overnight at 37 °C using trypsin (1: 40 w/w), Lys-C (1: 20 w/w), Lys-N (1: 50 w/w), Glu-C (1: 20 w/w), Asp-N (1: 40 w/w), chymotrypsin (1: 50 w/w) protease/substrate ratios. After enzymatic digestion, an equal volume of ethyl acetate was added, and the mixture was acidified with 0.5% trifluoroacetic acid (final concentration) according to the PTS protocol. The mixture was shaken for 1 min and centrifuged at 15,700 g for 2 min to separate ethyl acetate phase from the aqueous phase. The latter was collected and desalted by using SDB-StageTips.^44^ The amount of peptides was quantified by LC-UV at 214 nm relative to standard BSA tryptic digests and kept in 80% ACN and 0.5% TFA at −20 °C until use.

### Peptide fractionation by Strong Cation Exchange StageTip

The preparation of SCX-StageTips were performed in 200-μL tips format as described previously.^45^ SCX buffers were made in 15% acetonitrile with stepwise increase of elution buffer strength: F1 - 0.1% TFA; F2 - 1.0% TFA; F3 - 2.0% TFA; F4 - 3.0% TFA; F5 - 3.0% TFA and 100 mM AA; F6 - 3.0% TFA and 500 mM AA and; F7 - 0.1% TFA and 500 mM AA, as described previously.^46^ Two technical replicate SCX separations have been done for each digest. Conditioning and equilibration were done through sequential passing 100 μL buffer and centrifugation at 1000 × g for 1 min of the following buffers: methanol, F7, F5 and F1. 20 µg of digests from HeLa cell lysate were loaded into the SCX-StageTip, spun at 1000 × g for 1 min and the eluate was collected as flow-through (FT). The bound peptides eluted with 100 µL of F1 by centrifugation at 1000 × g for 1 min. Subsequent fractions were collected using 100 µL of SCX buffers F2 to F7. F5-F7 were lyophilized, resuspended in 50 μL of 0.1% TFA and desalted by SDB-StageTips.

### NanoLC/TIMS/Q/TOF analysis

NanoLC/MS/MS analyses were performed using a hybrid ESI/TIMS/Q/TOF mass spectrometer (timsTOF Pro, Bruker, Bremen, Germany), which was connected to an Ultimate 3000 pump (Thermo Fisher Scientific, Germering, Germany) and an HTC-PAL autosampler (CTC Analytics, Zwingen, Switzerland). Peptides were separated at 50 °C using 150 mm length × 100 μm ID capillary column with 6 μm ID ESI tip, packed with Reprosil-Pur 120 C18-AQ 3 μm particles (Dr. Maisch, Ammerbuch, Germany). The injection volume was 5 μL and the flow rate was 500 nL/min. The mobile phases consisted of (A) 0.5% acetic acid and (B) 0.5% acetic acid and 80% ACN. A two-step linear gradient of 5−40% B in 45 min, 40−99% B in 1 min, keeping at 99% B for 5 min was employed.

The timsTOF Pro mass spectrometer was operated in PASEF mode.^47^ Two methods were applied in IMS separation. Method 1 was applied for covering singly and multiply charged ions and method 2 was mainly used for depleting the contaminants usually singly charged background ions, respectively. The setting parameters are described in Table S1.

TIMS funnel’s voltages were linearly calibrated using Agilent ESI-L Tuning Mix to obtain reduced ion mobility coefficients (1/K_0_) for three selected ions (*m/z* 622, 922, 1222).^48^ The 1/K_0_ was converted to collisional cross section (CCS) using the Mason-Schamp equation.^49^

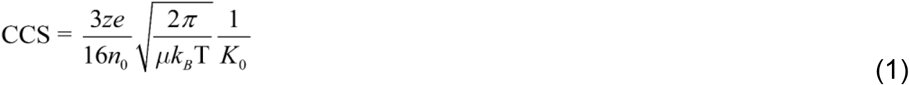

The z is the charge of the ions, e is the elemental charge (1.602 × 10^−19^ A·s), n_0_ is Loschmidt constant (2.686 × 1025 m^-3^), k_b_ is Boltzman’s constant (1.380 × 10^−23^ kg·m^2^·K^-1^·s^-2^), μ is the reduced mass (m_i_m_g_ / (m_i_ + m_g_), m_i_ is the mass of ion; m_g_ is the mass of N_2_, 1 Da = 1.660 × 10^−27^ kg), K_0_ is the reduced mobility, (10^−4^ cm^2^·V^-1^·s^-1^) and T is the temperature (305 K). For the CCS calculation, pure N_2_ is assumed as the drift gas.

### Peptide identification and retention time prediction data filtering

The peak list in mascot generic format (MGF) was generated by MaxQuant v1.6.7.0,^50^ encoding information on both retention time and 1/K_0_ for each spectrum. The peptides were identified using X!Tandem Cyclone (12.10.01.1)^51^ against human subset of the Swiss-Prot database (July 2016 extraction) with 20 ppm mass tolerance for both precursor and product ions. Carbamidomethyl (C) was set as a fixed modification. Oxidation (M, W), deamidation (N, Q), cyclization (N-term Q, C) and acetylation (protein N-term) were allowed as variable modifications, and strict enzymatic specificity allowing for up to 2 missed cleavages as search parameters. Redundant peptide identifications have been removed leaving the most intense peptide MS/MS hits with their correspondent 1/K_0_ and retention time values. Peptides with variable modifications were also removed for CCS prediction. All peptides with confidence score log (e) < -1 or better were additionally filtered using latest version of SSRCalc retention time prediction model.^39^ All peptides with retention time prediction error of more than ±6 min and low confidence score (−3 < log (e) <-1) have been removed as shown in Figure S1.

### Model optimization

The preliminary length-specific ensemble of multiple linear regression (LS-MLR) models by R package^52^ used to explore the variable space in CCS prediction has been derived for peptides with the selected charges and length (Eq.2):

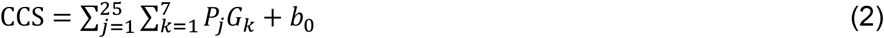

where, *P*_*j*_ is the position and group dependent coefficients, *G*_*k*_ is the mass of each amino acid and *b*_0_ is a constant. Amino acids have been grouped in seven categories based on their physicochemical properties as follows: basic K, R and H; acidic D and E; polar S, T, N and Q; aliphatic A, V, I and L; aromatic F, W and Y; aliphatic/polar side chains M and carbamidomethyl-Cys; P and G as amino acids with low helical propensity.

On the other hand, the final algorithm of the Sequence-Specific Ion Mobility Calculator (SSICalc) encodes 13 position-dependent ISP values (*j*) for each amino acid (*i*) in a charge (*z*) and protease (*e*) tryptic/non-tryptic dependence: six on each terminus plus internal position. Our equation for the SSICalc model is shown in Eq.3 as the summation of a coefficient (*P*) multiplied by the number of amino acids (*AA*) in the peptide with the corresponding *e, z, i, j* state listed above and mass of the amino acid (G_i_) along with a constant *b*_*0*_ term for the combined model:

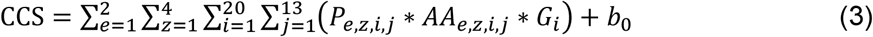

Optimization of the charge sub-divided models followed a simple stochastic hill-climbing approach maximizing to the highest R^2^ correlation. In each iteration of the optimization, a randomly selected parameter was adjusted along a shift value until the prediction versus observed CCS stopped improving until which a subsequent parameter is selected for optimization. The initial variable-space parameters were set to a matrix of ones and the signed shift value was randomly selected.

## Data availability

The MS raw data and analysis files have been deposited with the ProteomeXchange Consortium (http://proteomecentral.proteomexchange.org) via the jPOST partner repository (http://jpostdb.org)^53^ with the data set identifier PXD021440/JPST000959.

## RESULTS AND DISCUSSION

### Data selection for model optimization

In this work, seven protease-digested (trypsin, LysargiNase, Lys-C, Lys-N, Glu-C, Asp-N and chymotrypsin) cell lysates have been analysed using SCX-StageTip fractionation applied prior to RP-LC/TIMS/Q/TOF analysis. SCX chromatography was chosen aiming to improve representation of peptides in different charge states. The selection of peptides for model optimization are crucial for generating a representative high-quality dataset. NanoLC/TIMS/Q/TOF measurements provided the reduced ion mobility coefficients (1/K_0_) and retention time for all identified peptides. These identifications have been additionally filtered using the SSRCalc peptide retention time prediction model as shown in Figure S1. Less than 1% of identified peptides were removed based on large retention prediction errors or low confidence score of -3 < log(e) < -1. Moreover, since IMS separates peptides based on their conformations similar to previous studies,^32,34^ multiple peptide conformations were detected in some instances (Figure S2). The 1/K_0_ values for model optimization were then selected corresponding to the most intense peptide MS/MS hit on each mobilogram (Figure S2) followed by the removal of redundant identifications in order to merge the dataset into 133,946 entries. There were 14,482, 86,268, 27,463 and 5,733 peptides belonging to the 1+, 2+, 3+ and 4+ populations, respectively. Peptides contained 1-11 positively charged residues (Lys, Arg, His and unmodified N-termini) and were 5-50 amino acids long (560-5245 Da): 14.7 residues on average. This represents a typical size/charge distribution of peptides encountered in bottom-up proteomics experiments.

### Evaluation of peptide bulk properties affecting CCS

Similar to prior work,^32,35^ plotting the dependence of CCS values on *m/z* resulted in definitive trend lines corresponding to four individual charges (Figures 1, 2). We used the characteristic shapes of these plots and properties of 100 peptides that are the most significant positive/negative outliers (Table 1, Figure S3) for each charge state to assess the effects of peptide bulk properties.

**Table 1.** Average bulk properties of top 100 positive and negative outliers in charge specific CCS vs. *m/z* plots.

| Charge/<br>prediction<br>error | Tryptic/non-tryptic<br>peptides* | Peptide<br>length | Agadir α-<br>helicity<sup>54</sup> | SSRCalc<br>Hydrophobicity<br>(% ACN) | pI<sup>55</sup> | # of basic<br>residues |
| --- | --- | --- | --- | --- | --- | --- |
| 1+/pos | 93/7 | 12.60 | 0.31 | 11.88 | 7.13 | 2.00 |
| 1+/neg | 6/94 | 13.10 | 0.11 | 7.92 | 3.55 | 1.23 |
| 2+/pos | 70/30 | 18.56 | 1.41 | 15.35 | 6.88 | 2.93 |
| 2+/neg | 44/56 | 21.52 | 0.16 | 12.09 | 4.44 | 2.19 |
| 3+/pos | 60/40 | 29.44 | 1.35 | 18.43 | 5.97 | 3.41 |
| 3+/neg | 27/73 | 28.87 | 0.23 | 12.96 | 4.30 | 2.81 |
| 4+/pos | 42/58 | 33.77 | 0.68 | 16.01 | 6.42 | 4.34 |
| 4+/neg | 40/60 | 36.27 | 0.32 | 15.67 | 4.44 | 4.12 |

\* - independently of the protease used tryptic peptides correspond to Lys/Arg terminated species.

**Figure 1.**
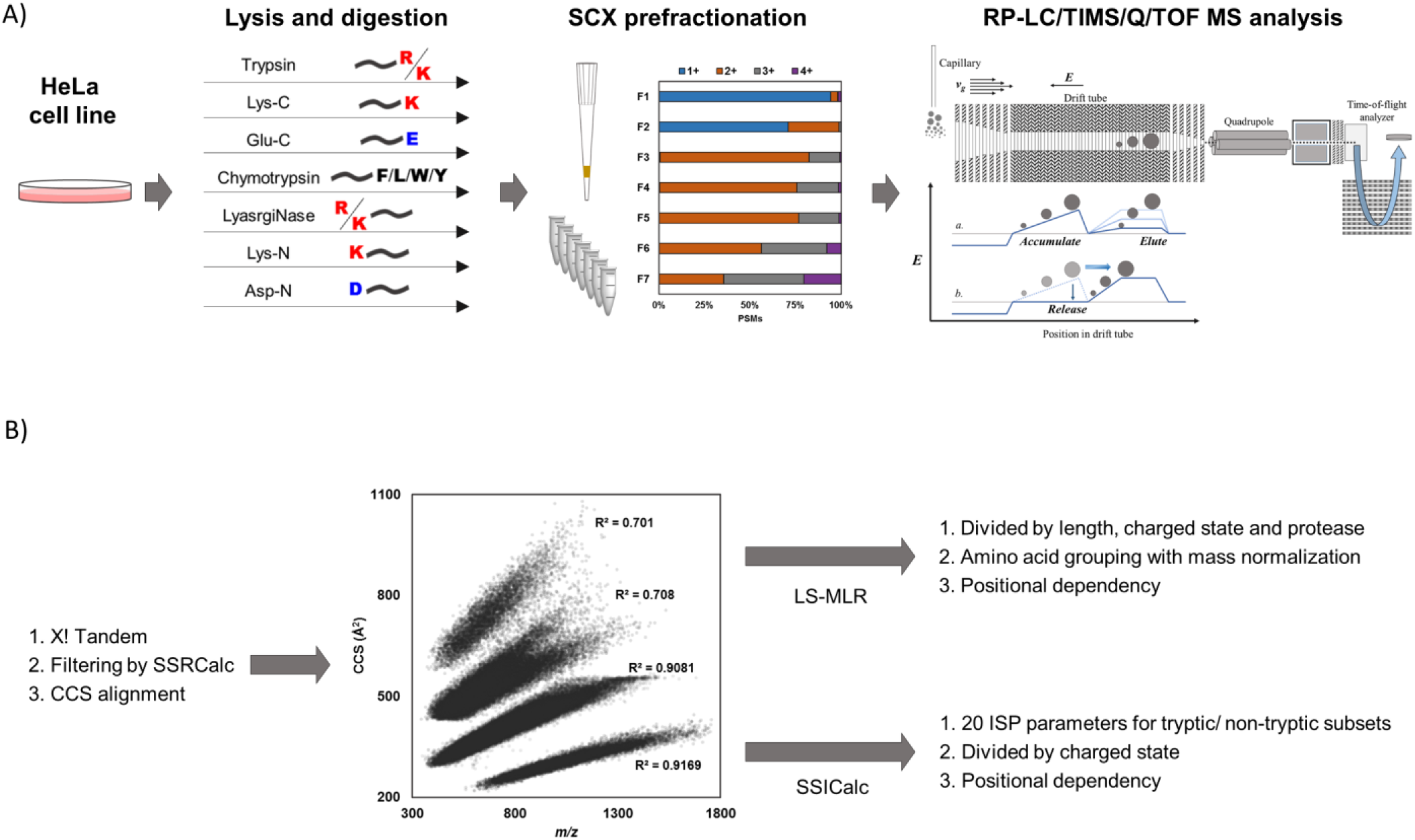
An overview of the workflow used in our work. (A) Experimental data collection, (B) prediction model optimization.

**Figure 2.**
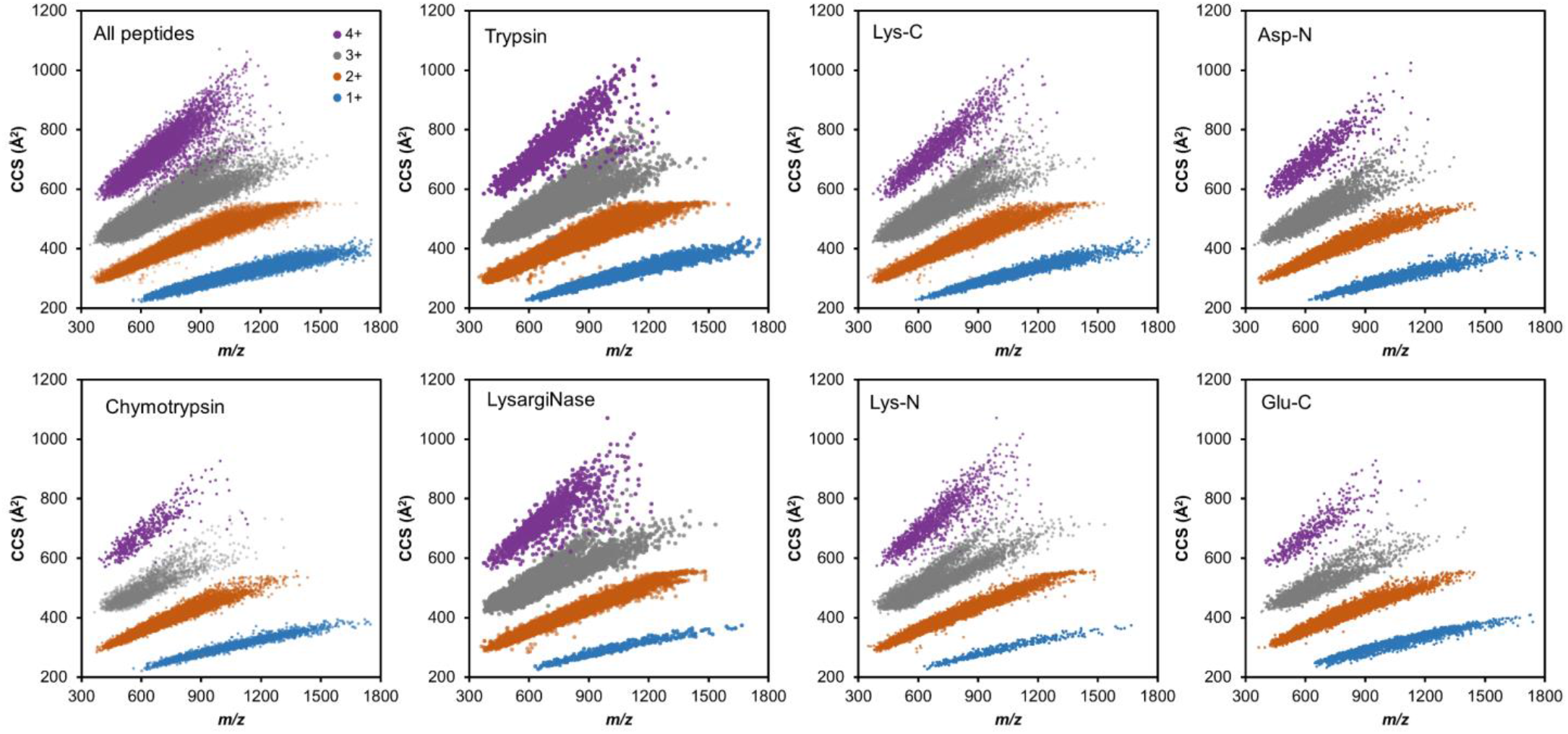
CCS versus *m/z* plots for 133,946 peptides from HeLa cell digests for total (A) and protease-specific populations (B-H). Individual charge states are color coded as: blue, 1+; orange, 2+; gray, 3+ and purple, 4+.

The CCS versus *m/z* correlation plots for singly and doubly charged peptides are slightly concave, indicating the preference of longer peptides to be in more compact conformation. For longer highly charged (3+, 4+) peptides, the CCS trends became dispersed and a clear split-population appeared in 3+ species, corresponding to compact (low CCS) and extended (high CCS) features observed previously.^32,34,35^ In addition, we found that the distributions between compact and extended conformation is protease-specific, e.g., 3+ Lys-N digested peptides containing two positive charges at N-termini predominantly assume compact conformation (Figure 2). The non-tryptic (terminated by any amino acid other than Lys or Arg) peptide populations of all charges exhibited lower CCS values compared to tryptic ones. This observation was also confirmed by analyzing the population of the outliers (Table 1). For example, 93% of 1+ peptides with largest positive deviations are tryptic, while 94% non-tryptic species were found among 100 most negative outliers. Similar finding has been reported by Lietz et al., who compared CCS values for 3+ peptides from Lys-C and Lys-N digest. The authors explained this behaviour by the electrostatic interaction of N-terminal/C-terminal Lys with peptide macro-dipole, which should destabilize/stabilize peptide’s helical conformation, respectively. Similarly, 3+ Asp-N peptides exhibit even distribution between compact and extended structures, while Glu-C generated species tend to be in the latter conformation (Figure 2). Overall, LysargiNase/Lys-N/Glu-C destabilize the helix favouring compact, whereas trypsin/Lys-C/Asp-N stabilize the helix inducing more extended form through interaction of terminal residues with peptide’s macro-dipole.^34^ Detailed analysis of 1+ and 2+ correlation plots (Figure 2) also shows that trypsin/Lys-C/Asp-N populations show some splitting between the dominant compact and extended subpopulations, although the CCS difference between two conformations was subtle. Similarly, LysargiNase/Lys-N/ Glu-C exhibit more uniform distribution in the 1+ and 2+ peptide populations. Furthermore, for 4+ trypsin/Lys-C/Asp-N, more preference to the extended conformation was observed compared to LysargiNase/Lys-N/Glu-C. For each protease, increasing charge state of a peptide led to higher tendency to be in extended conformation.

Based on comparison of average peptide length for the entire population (Table S2) vs. the most significant outliers (Table 1), we find that shorter peptides are more consistent in conformational behaviour than the longer outlier peptides. In other words, shorter peptides are characterized by smaller prediction errors. Longer peptides have more accessible conformational states that can be either more compact or extended than the average population. This behaviour is particularly obvious for 3+ peptides, which exhibit a split for more than 20 mer species. Peptides with large positive prediction errors exhibit higher helical content calculated using the Agadir algorithm.^54^ Alpha-helical peptides are more linear in geometry and are unable to fold to smaller states thus will exhibit higher than expected CCS consistent with the positive prediction error we observed. Peptides with large positive deviations (Table 1) are more hydrophobic. This observation supports previous findings by Valentine et al.^31^ and Shvartsburg et al.^56^ reporting on high ISP values of hydrophobic residues.

The peptide pI showed no obvious correlation with deviation from CCS vs. *m/z* plots, when entire peptide population was considered (Figure S4). However, when we isolate the top 100 outliers in the dataset, as shown in Table 1, the general trends amongst all the charges are that positive prediction errors are associated with higher average pI values and vice versa. We have an interest to investigate if there is a correlation between peptide CCS and electrophoretic mobility measured by capillary zone electrophoresis (CZE). Our advanced SSRCalc CZE model has R^2^ ∼ 0.995 correlation with experimental values^42^ and should provide an accurate estimation of electrophoretic mobility in solution at acidic pH when compared to experimental CCS. However, Figure S5A shows poor correlation between these two systems. Important to note that this plot consists of multiple sub-populations corresponding to peptides carrying different number of charged residues versus their CCS values for a particular charge state. Peptide electrophoretic mobility at acidic pH depends mostly from peptide charge (number of basic residues) and mass. The sequence-specific features in CZE largely are limited by N-terminal position of Asp and Glu, which reduces N-terminal charge/basicity. IMS separation are affected by many processes, including formation of helical structures, which results in poor correlation of CCS versus µ_ef_ plots even when peptides with identical number of charged residues are considered. As shown previously,^42^ the semi-empirical model µ_ef_ = k(Z/M^X^) can be optimized by modulating X such as 1/3, 2/3, or 1/2. As CCS is proportional to mass (in Figure 1), we were able to linearize the CCS versus µ_ef_ correlation by modulating X to 1/3 with R^2^ = 0.849 (Figure S5B).

### Length-specific multiple linear regression model

To explore the properties involved in the ion mobility of peptides in IMS, an ensemble of length-specific multiple linear regression (LS-MLR) model and charge states was built to predict the CCS values of the peptides. In each model, the number of independent variables increases with peptide length, as there are 20 amino acids per position in the peptide sequence. However, due to the nature of enzymatic digestions, the number of experimental data per peptide length was only distributed over a narrow range, and in particular, the number of longer peptides was sparse (Figure S6A). To reduce the number of independent variables, in addition to the terminal cleaved site, the 20 amino acids were grouped into 7 by similarity and LS-MLR analyses were performed using the relative position coefficients (*P*_*j*_) corrected for the mass (*G*_*k*_) of each amino acid (Figure 1). Different proteases cleaved at N/C-terminal sides of different amino acids, resulting in the diverse CCS values observed in IMS (Figure 2). Therefore, individual LS-MLR models were built for peptides generated by different proteases and in different charged states, and the position coefficients were converted to coefficients for each amino acid based on mass as shown in Figure 3.

**Figure 3.**
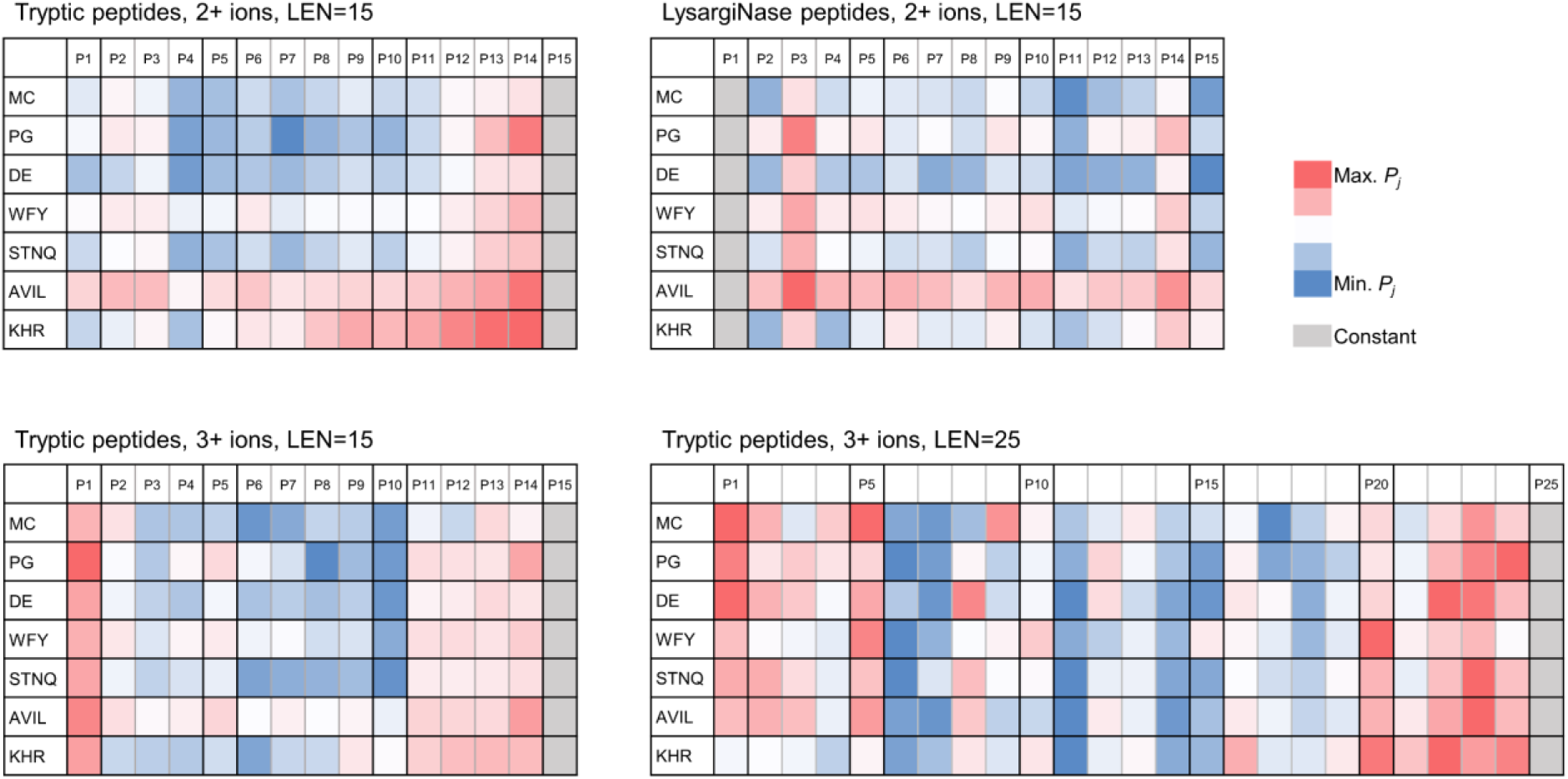
Heatmaps of position and group dependent coefficients obtained by LS-MLR model.

As a first step, LS-MLR was applied to trypsin- and LysargiNase-digested peptides that have opposing terminal Lys/Arg (Figure 3). The trypsin-digested peptides have higher coefficients at the C-termini especially for aliphatic and positively charged amino acids, whereas positive contribution at P3 position was observed for LysargiNase-digested peptides. Second, both 2+ and 3+ tryptic had larger coefficients at the C-termini, and only the 3+ peptides had larger coefficients at the N-termini. Moreover, some polar amino acids such as D, E, S, T, N, and Q have lower coefficients in the internal region. Finally, LS-MLR was performed for each length of tryptic peptides. The bottom two heatmaps (Figure 3C and 3D) show the coefficients in CCS prediction for 3+ tryptic peptides of different lengths. At a peptide length of 15, the coefficients are higher for both termini and lower for the polar amino acids in the central region. However, as the length of the peptide increased, the coefficients become more uniform in the central region.

The LS-MLR model has been able to achieve an R^2^ value of 0.977 for the CCS prediction derived from the tryptic peptides for specific charge and length (Figure S6B, 1+ peptides with 7-12 a.a., 2+ peptides with 8 -20 a.a. and 3+ peptides with 11-25 a.a.). While the LS-MLR models could produce good correlations for peptides of particular length, the overfitting still occurred due to the limited size of the dataset at each length. Therefore, it is necessary to apply an alternative predictive model that is not limited by the number of features to obtain a global prediction of CCS values. To compensate for these features, we employed the physicochemical properties of trypsin/non-trypsin peptides, charge states, and amino acid positions via a step-by-step charge and protease dependent ensemble linear regression model optimization, as shown in the next section.

### Peptide length independent step-by-step optimization using Intrinsic Size Parameters (ISP) approach as a starting point

Each optimization step has been followed by the alignment of eight peptide subsets: tryptic/non-tryptic in four different charge states to fit all eight correlation plots to CCS_pred_ versus CCS_exp_ with slope 1 and intercept 0 as shown in Table S2. In every step each of the eight independently optimized sub-models have their own CCS-aligning slope and intercept values that are allowed to vary from this initial *m/z* data alignment. It should be noted that the R^2^ correlation for the combined collection of peptides is higher than individual subgroups due to wider range of the experimental CCS values.

Step 1: twenty ISP values optimized for two datasets (tryptic and non-tryptic). Figure 4A shows comparison of ISPs reported by Valentine et al.^8^ and by Shvartsburg et al.^56^ versus ours optimized for tryptic/non-tryptic datasets. Most of the hydrophobic residues’ ISP values are larger compared to the ones reported previously. Conversely, polar residues showed lower ISP values, favouring more compact structures. These deviations are likely originated from the significant difference in charge and size distribution in these two datasets. ISP values for Lys and Arg, which have been found to be similar to His and in close agreement with a-priori predicted ISP values by Shvartsburg et al. using sum of the projection areas for constituent atoms.^56^

**Figure 4.**
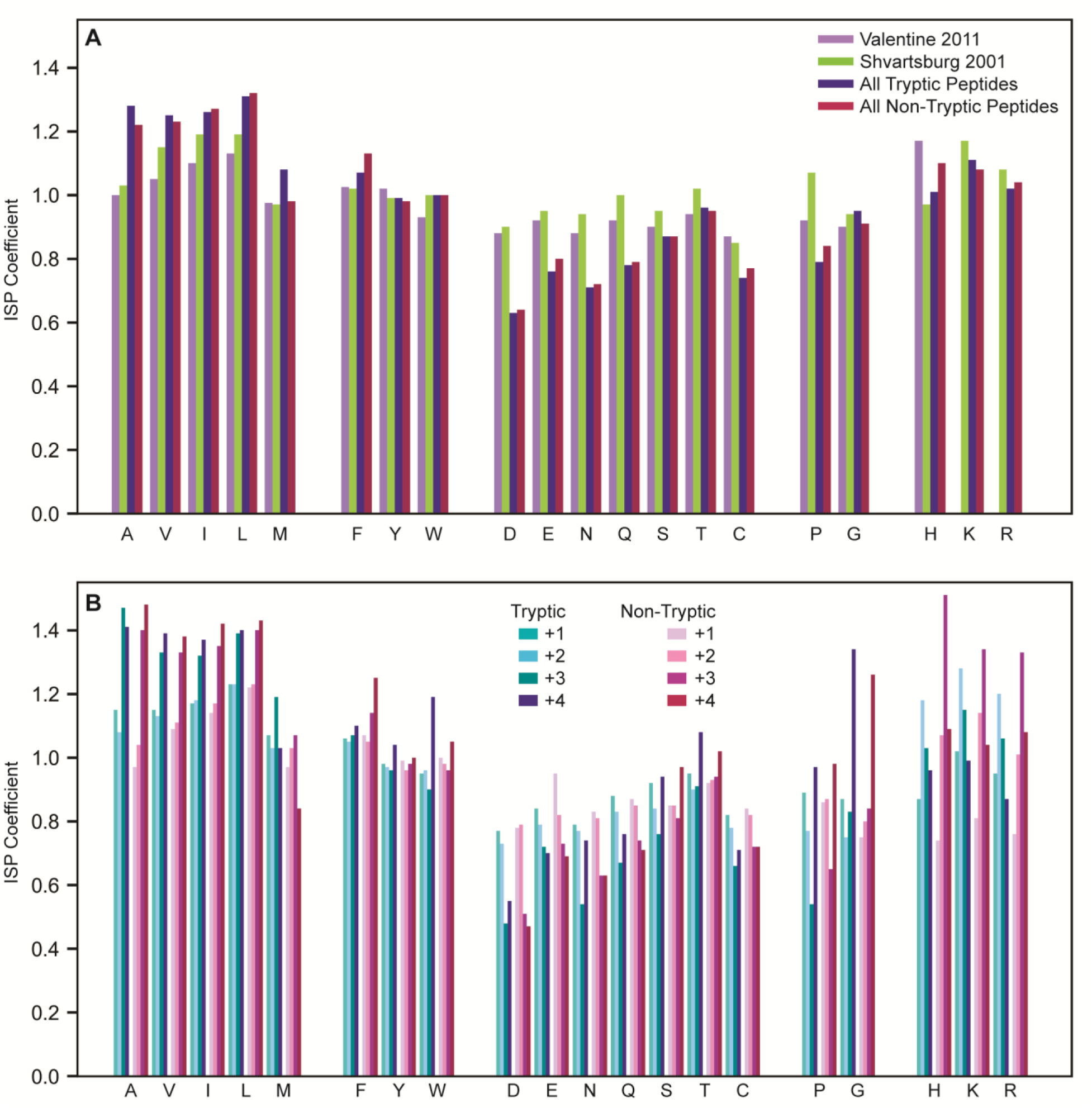
Comparison of previously reported ISP values with ones obtained for our dataset. A – ISP values for tryptic and non-tryptic peptides vs. Valentine at al.^8^ and Shvartsburg et al.^56^ data (Model Step 1); B – charge specific ISP values (Step 2).

Step 2: charge dependent ISP values have been optimized for subsets of peptides and improved correlation values for all of the submodels (Table S2). Figure 4B shows these values for both tryptic and non-tryptic species. ISP values of hydrophobic residues increase for highly charged 3+ and 4+ peptides; the opposite is true for polar residues (D, E, N, Q). These trends follow the difference observed between Valentine et al. values and ours, indicating inclusion of highly charged longer peptides determined overall differences in ISP values in Figure 4A. Pro exhibits the lowest ISP values, favouring compact structures, for 3+ peptides, while Gly in 4+ species promote extended conformation.

Step 3: thirteen position dependent ISP coefficients have been introduced for each residue: six on both termini plus a general internal position – similar to all SSRCalc models for peptide HPLC. This led to further improvements in correlation values in all respective subsets shown in Table S2. Selected position dependent trends are shown in Figure 5 and the entire collection of optimized coefficients is provided in Table S3.

**Figure 5.**
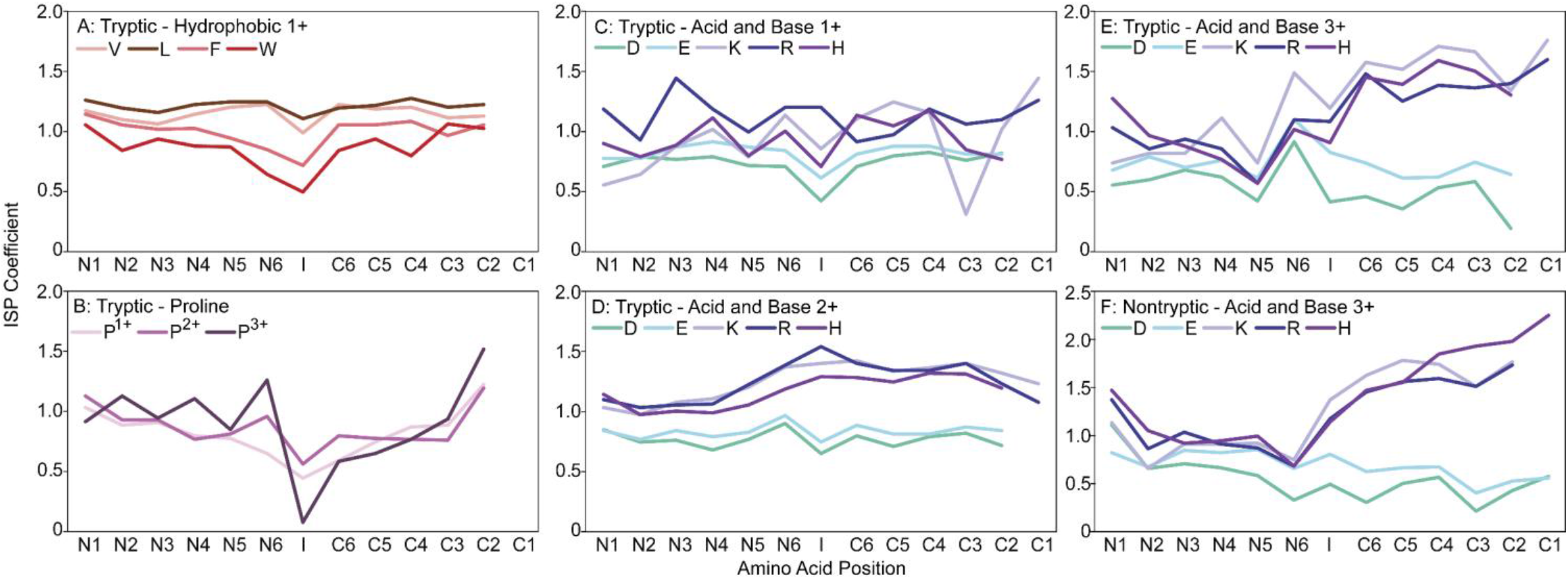
Selected examples of ISP positional trends: for hydrophobic amino acids (A), Pro in different charged peptides (B), acidic/basic amino acids among different charged peptides (C, D, E, F).

The hydrophobic residues (A, V, L, I, M, F, Y, W) show virtually no position dependence except for a small decrease in ISP for internal position, especially for aromatic Phe, Tyr, and Trp (Figure 5A). Also, an evident decrease in the internal position for Pro. It is interesting to note that Pro ISP values in terminal positions are above 1, which corresponds to the value determined by Shvartsburg et al. based on atom projections.^56^ In other words, the behavior of Pro (low ISP) is determined by its known property as helix breaker, rather than size of the side chain. While located inside of the peptide Pro tends to form kinks in the structure favouring sequence bending, resulting in reduced CCS. The decrease in internal Pro ISP is smaller for 2+ ions compared to 1+ and much more significant for 3+ (Figure 5B).

The position dependence for basic Arg, His, Lys is also unique (Figure 5C-F). Generally, ISP values increase slightly from N- to C-terminus corresponding to their increasing interactions with the helix macro-dipole near C-terminus. This trend is more visible for 2+ and 3+ peptides compared to 1+. Polar acidic residues exhibit lower CCS values for 1+ internal position (Figure 5C), which is similar to Pro. Asp, and Glu ISP values for 3+ peptides showed the effect of interaction with macro-dipole opposite to the basic residues as ISPs decrease from N to C-terminus.

While we do have a diverse set of peptides derived from different protease cleavages, there could be some issues with representation of amino acids at particular positions which will result in model over-fitting. For example, position dependent ISP values for charged residues showed significant variation due to their small representation in the 1+ subset. Position dependent ISP values for 4+ charge state also vary significantly, making it hard to extract consistent ISP trends.

In other words, CCS prediction model for the 5,733 4+ peptides with 520 parameters is over-fitted. Additive retention time prediction models in peptide RP-HPLC show representative results starting at a 1:5 parameter to peptide ratio,^57^ suggesting significant variation in peptide conformation in IMS separation. Nonetheless, overall model was able to achieve an R^2^ value of 0.981 and demonstrated robustness in predicting CCS values consistent with experimental trends.

### Composition and sequence-specific features driving peptide IMS separation

The original work on incorporating ISP concept has been done using collections of structurally similar 1+ and 2+ tryptic peptides without internal Arg/Lys residues.^31^ The largest dataset used by Valentine et al. consisted of 2,094 peptides, 10.7 residues long on average.^8^ We anticipated that inclusion of the entire population of peptides without restriction on protease type, number of basic residues, charges and peptide length will complicate model optimization. At the same time, it has provided additional information on the mechanism of ion mobility separation. Due to the increased size of the dataset, we were able to elucidate position-dependent ISP and found significant effect of the structural features rather than geometric size of individual residues.

The geometry of peptides in gas phase are strongly affected by the charge of the peptides. As seen in the plot of CCS versus molecular weight (Figure 6A), increasing in peptide charge leads to higher CCS values. To explain our findings, we use Counterman & Clemmer’s approach^33^ that have described the notion of exposed cationic charges being solvated by the backbone carbonyls of the peptide leading to the compact globular structures (Figure 6B). From our results in proteomic IMS separations, it suggests that ability of a peptide to solvate the charge as a globular peptide is based on the peptide flexibility and availability of polar groups as elaborated below in the instances of 1+ peptides. As charge density increases, the repulsive effects of cationic charges in proximity starts to be become an issue in charge solvation. The repulsion reduces the stability of globular structures and starts to approach other stable conformations that are more linear in orientation. The different stages of peptide structures will be described as: closed globular, open globular, hinged-helix, alpha-helix, and linear listed in order of increasing CCS.

**Figure 6.**
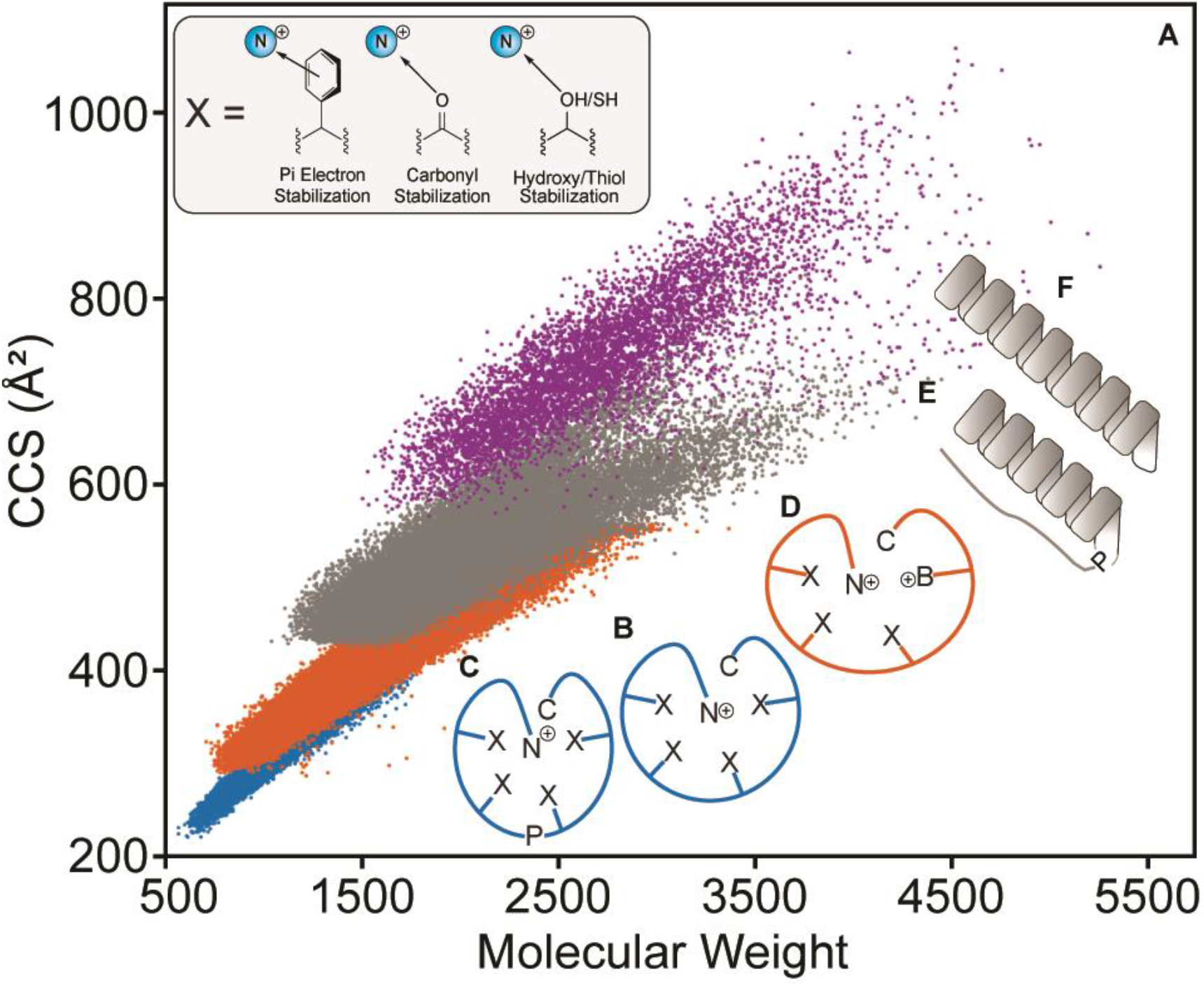
Compositional and sequence specific features driving the separatory behaviour observed in CCS vs. Molecular Weight plot (A). The geometry of the peptides in gas phase (C, B, D, E, F) ordered from lowest to highest CCS. N^+^ corresponds to the N-terminal, C to the C-terminal, B^+^ is an internal basic residue, and P is Pro.

In singly charged peptides (1+), the predominant geometry will be closed globular allowing this group to have the lowest CCS. The amino acid side chain structure will influence the size of the globular peptide based on steric interactions and electronic effects. The solvation of the exposed cation will be enhanced in the presence of partially negative functional groups as shown in Figure 6B. Acidic Asp and Glu show low ISP coefficients as they improve the solvation with their carboxylate side chains, which assist in compacting the globular structure. Asn and Gln also follow a similar trend where the carbonyl on the amide also assist in compacting the globule. Cys, Ser and Thr have polar thiol or hydroxyl groups that can stabilize the cation therefore exhibiting low ISPs. Aromatic amino acids stabilize the peptide electronically via the pi orbitals on the aromatic rings and are able to condense the peptide structure. In the case of aliphatic amino acids, their side groups do not contribute electronically to the cation stabilization but rather add steric bulk to the peptide leading to the observed increase in CCS. The flexible Gly and Pro do not contribute electronically or sterically but rather their flexibility allows for tighter peptide solvation to the cationic core allowing lower CCS conformations (Figure 6C).

For the case of doubly charged peptides (2+), they follow a similar trend in internal ISP values with +1 peptides; however, exhibiting smaller change in ISP values between terminal and internal positions. This suggests the 2+ peptides are also globular but given the electrostatic repulsion between the two positive charges, the peptide will not be able to fold as tightly (Figure 6D). This effect forces 2+ peptides to the open globular conformation consistent with the higher CCS than 1+ peptides found in our experimental data. This observation is supported by the divergence in acidic and basic amino acid ISP values shown for 1+ and 2+ peptides in Figure 5C and D where the mediation of an acidic side chain assists in lowering the repulsive effects therefore decreasing CCS and vice versa for basic amino acids.

The triply charged peptides (3+) exhibit a divergent pattern when CCS is plotted against molecular weight. Prior work in the field has demonstrated that the pattern can be attributed to two main peptide geometries:^33^ a fast hinged-helix orientation and a slower alpha-helix orientation as displayed in Figure 6E, F. As peptides are now able to uptake helical conformations in this charge state, the ISP values of aliphatic amino acids are increased from their 2+ counter parts. The helix-breaker Pro exhibits the lowest ISP value for +3 charge state (especially for internal positions as shown in Figure 5B) due to their ability to bend the peptide to favour the hinged-helix orientation allowing the peptide to have lower CCS. Similarly, acidic and polar amino acids also decrease in CCS from 2+ peptides as the effect of the cationic solvation is more drastic in larger ions found in the 3+ sets (Figure 4B).

Our findings on splitting population of 3+ peptides were confirmed by protease-specific features of CCS vs. *m/z* plots driven by interaction of acidic/basic residues with peptide macro-dipole. Peptides featuring acidic residues at C-termini and basic ones at N-termini (LysargiNase/Lys-N/Glu-C) tend to be in a compact conformation. Meanwhile trypsin/Lys-C/Asp-N peptides show more even distribution between two conformational states. Surprisingly, ions in the other charged states (1+, 2+ and 4+) also showed similar specificity, albeit with uneven distribution between conformational states. Lesser number of 1+ and 2+ peptides assume extended and 4+ compact conformation, respectively. Compared to previous studies, we can identify this novel finding due to the diversity of proteases employed.

Quadruply charged peptides (4+) in the past have not been well characterized due to their limited representation in the optimization datasets.^58^ Based on our novel CCS information we conclude that the geometry of the peptides are generally more linear and helical than 3+ peptides. In terms of the aliphatic amino acids, the ISP values are largely similar to 3+ peptides (Figure 4B) supporting our notion that helicity is still a strong contributor in the 4+ charge state. Interestingly, Pro and Gly increase in ISP values (Figure 4B). Electrostatic contributions from acidic amino acids in quadruply charged peptides are analogous to the triply charged peptides, whereas basic amino acids experience a decrease in ISP values (Figure 4B).

## CONCLUSIONS

Through pairing high-throughput proteomics with IMS, we were able to collect a high-quality CCS database of ∼134,000 peptides and establish first sequence-specific model to predict peptide CCS. Our collections and resultant prediction model are detailed for each charge state, in enabling expansion for the current observations of 3+ and 4+ peptides in finer detail, and in attaining an R^2^ value of 0.981 for the entire dataset. The gas phase peptide geometry dictates the CCS of the peptide and the conformations are heavily influenced by charge, sterics, and helical propensity of the constituent amino acids. Singly charged peptides have the lowest CCS in the entire dataset as it maintains a small profile in a closed globular conformation with the cation stabilized by backbone carbonyls or polar side groups. The globular structure can be further stabilized and condensed in the presence of Pro. For doubly charged peptides, the geometric behaviours are similar to 1+ peptides; however, the two cations experience electrostatic repulsion causing the structure to expand to an open globular conformation. Triply charged peptides establish two main conformations, a fast hinged-helix or a slow alpha helix structure. The ISP contributions of hydrophobic amino acids increase compared to the previous two charge states as these amino acids have high helical propensity favouring the alpha-helix conformation. Pro also exhibits the lowest ISP in 3+ peptides as its ability to bend the peptide favour the formation of the hinged-helix structure. We observe a divergent trend between acidic and basic amino acids’ position dependent ISPs in triply charged peptides due to the macro-dipole interaction, which is also characteristic for helical structures. For the first time, 2+ peptides as well as 1+ and 4+ peptides were identified to exhibit similar splitting behaviour, due to the position of acidic/basic residues that favour helical stabilization via interaction with the peptide macro-dipole. Quadruply charged peptides maintain similar ISP values and trends as 3+ peptides with the exception of Pro and Gly increasing drastically in ISP. Other structural outliers have been observed for long and highly charged peptides with multiple proline residues. These motifs are one of the reasons for the high prediction errors observed in 3+ and 4+ peptides as the interactions of adjacent prolines may result in the formation of left-hand helices, which extends the peptide conformation. There is an active effort to understand the behaviour of different polyproline isomers; however, with current literature it is difficult to definitively align our diverse observations for such species. To fully elucidate the nature of our prediction errors, molecular dynamics paired with hydrogen-deuterium exchange experiments for a majority of the peptides will be needed to understand the true diversity of gas-phase peptide conformations. Despite the difficulties in ascertaining outlier behaviours in our dataset, we are able to provide a variety of novel insights for the influence of peptide properties in real world CCS prediction.

## Supporting Information

Figure S1: Elimination of false-positive identifications using peptide retention time prediction with de-novo SSRCalc model (A, B) or SSRCalc retention Database (C, D). All peptides with retention time prediction error of more than ±6 min and low confidence identification scores (−3 < log (e) <-1) have been removed.

Figure S2: Distribution of extracted ion (754.04-754.06 *m/z*) in chromatogram and mobilogram. The gradient blue on the right shows the intensity scale in MS1, retention time on y-axis, and 1/K0 on x-axis. Extracted ion chromatogram and extracted ion mobilogram are projected on the left and bottom axes, respectively.

Figure S3: Graphical representation of the selection process for the Top 100 Positive Outliers and Top 100 Negative Outliers shown in Table 1. The outlier selection lines (red) are parallel to the trendline of the CCS vs *m/z* correlation (black dotted) as we are picking the outliers with the largest positive difference for the Top 100 Positive Outlier set and the largest negative difference for the Top 100 Negative Outlier set.

Figure S4: Correlation between CCS prediction error and peptide pI.^55^ Prediction errors have been calculated for uncorrected CCS vs. *m/z* plots for each charge state.

Figure S5: (A) Correlation between predicted peptide electrophoretic mobility^42^ and experimental CCS values. Peptides detected in 2+ charge state by mass spectrometer, but carrying different charge (2+, 3+, 4+, 5+) in solution at acidic pH are highlighted. (B) As shown in the paper,^2^ the semi-empirical model µ_ef_ = k(Z/M^X^) can be optimized by modulating X such as 1/3, 2/3, or 1/2. As CCS ∝ M (in Figure 6), we attempted the X modulation and 1/3 had the best correlation of R^2^=0.849.

Figure S6: (A) The number of peptides for each charge state and length in tryptic dataset compared to the number of features for each length in the LS-MLR. (B) Correlation of predicted versus experimental CCS values. The predicted CCS values derived from the tryptic peptides for specific charges and length (1+ peptides with 7-12 a.a., 2+ peptides with 8 -20 a.a. and 3+ peptides with 11-25 a.a.) by LS-MLR.

Table S1: Parameter settings of timsTOF Pro mass spectrometer in PASEF analysis.

Table S2: Accuracy of prediction model (R^2^-value) for step-by-step optimization.

Table S3: Intrinsic size parameter values for step-by-step optimization of SSICalc model.

## ACKNOWLEDGEMENTS

This work was supported by the grants from JST Strategic Basic Research Program CREST (No. 18070870), JSPS Grants-in-Aid for Scientific Research (No. 17H05667), and the Natural Sciences and Engineering Research Council of Canada (RGPIN-2016-05963; O.V.K.). The authors also thank Dr. L. P. Kozlowski for assistance with isoelectric point calculations and Ryo Kajita (Bruker Japan K.K.) for technical support.

